# KDM3A regulates alternative splicing of cell-cycle genes following DNA damage

**DOI:** 10.1101/2020.02.26.965970

**Authors:** Mai Baker, Mayra Petasny, Mercedes Bentata, Gillian Kay, Eden Engal, Yuval Nevo, Ahmad Siam, Sara Dahan, Maayan Salton

## Abstract

Changes in the cellular environment result in chromatin structure alteration, which in turn regulates gene expression. To learn about the effect of the cellular environment on the transcriptome, we studied the H3K9 de-methylase KDM3A. Using RNA-seq, we found that KDM3A regulates the transcription and alternative splicing of genes associated with cell cycle and DNA damage. We showed that KDM3A undergoes phosphorylation by PKA at serine 265 following DNA damage, and that the phosphorylation is important for a proper cell cycle regulation. We demonstrated that SAT1 alternative splicing, regulated by KDM3A, plays a role in cell cycle regulation. Furthermore we found that KDM3A’s demethylase activity is not needed for SAT1 alternative splicing regulation. In addition, we identified KDM3A’s protein partner ARID1A, the SWI/SNF subunit, and SRSF3 as regulators of SAT1 alternative splicing and showed that KDM3A is essential for SRSF3 binding to SAT1 pre-mRNA. These results suggest that KDM3A serves as a sensor of the environment and an adaptor for splicing factor binding. Our work reveals chromatin sensing of the environment in the regulation of alternative splicing.

## INTRODUCTION

Splicing of precursor mRNA (pre-mRNA) is an important regulatory step in gene expression. The process of alternative splicing enables the production of protein isoforms from a single gene, isoforms that can have different functions, all contributing to the cell’s diverse tasks^1^. Traditionally the regulation of alternative splicing was attributed to regulatory elements in the pre-mRNA. Recently we and others have shown the link between chromatin structure and epigenetic modifications in splicing regulation^2-10^. The data linking chromatin and alternative splicing is not enough to explain the physiological causes and consequences of chromatin-mediated changes in alternative splicing. Furthermore, a major unanswered question in the field is how, and to what extent, environmental changes shape the cell transcriptome via modulation of alternative splicing.

Cellular environment has been shown to modulate splicing in several ways^11,12^. The main one is the induction of post-translational modifications of splicing factors. Phosphorylation of SR splicing factors affects their protein-protein interactions as well as regulating protein activity^11^. A specific example is activation of the Fas receptor leading to dephosphorylation of SR proteins that promote pro-apoptotic isoforms of two key regulators, Bcl-X and caspase-9^13^. Another example is cell-cycle-dependent splicing patterns, which are also modulated by SR protein phosphorylation by specific kinases and phosphatases such as NIPP1^14^. Another mode of action for alternative splicing regulation following cellular environmental changes could be by modulating chromatin, as changes in the environment are sensed by cells and transduced into changes in gene expression^15^, leading to specific cellular responses.

The histone demethylase KDM3A has been shown to respond to the environment to alter transcription patterns^16,17^. KDM3A’s specific demethylase activity is on H3K9me1 and H3K9me2 catalyzed by histone methyltransferases EHMT2/G9a and EHMT1/GLP^18,19^. Unlike H3K9me3, which is mostly a repressive mark present in heterochromatin regions, H3K9me2 is mainly on euchromatin^18,19^. The regulation of H3K9me2 is dynamic and KDM3A is known to be regulated post-translationally in multiple ways^16,17^ and thus it is suitable to be a responsive sensor for the environment.

KDM3A was shown to be phosphorylated in response to cold temperatures in adipose tissue and promote a specific expression pattern by demethylating promoters of genes important for chronic adaptation to cold stress^16^. In addition to cold shock, KDM3A is also phosphorylated on serine 265 following β-adrenergic stimulation. Interestingly, the modification promotes a scaffold function of KDM3A separate to its demethylase activity^17^. Serving as a scaffold for acetyltransferases present in enhancers and promoters allows transcription of key metabolic genes^17^. Under heat shock conditions phosphorylation of KDM3A on serine 264 promotes transcription of specific target genes^20^. Another response by KDM3A is to pressure-overload by cardiomyocytes which promotes transcription of genes with pro-fibrotic activity^21^. In addition, KDM3A responds to IL-6, androgen receptor and hypoxia^22-24^.

KDM3A’s pivotal role in transcription suggests that its misregulation could be a cause of disease. Indeed upregulation of KDM3A is related to tumorigenesis of colorectal cancer cell migration and invasion^25^. This function of KDM3A is attributed to its transcription of c-*Myc*, cyclin D1, and *PCNA*^26,27^. In addition regulation of androgen receptor and estrogen receptor transcription by KDM3A position it as a central player in prostate and breast cancer ^23,28-32^.

The role of H3K9me2 modification in alternative splicing was demonstrated by us and others. While H3K9me2 modification in promoter regions have been shown to cause gene silencing, the same modification in the intragenic region of genes was shown to alter alternative splicing^28,33-41^. The connection of the H3K9 methylation and the spliceosome was shown to be via an adaptor system containing a chromatin reader, which is one of the three HP1 proteins known to bind H3K9 methylation, HP1α, β and γ^42^. The chromatin reader in turn will bind a splicing factor that will recruit or block the spliceosome to promote a specific event of alternative splicing^33^.

Here we explored KDM3A’s role in the cellular response to the changing environment. To this end we conducted RNA-seq following silencing of KDM3A and discovered a functional group of cell-cycle genes to be regulated in both expression and alternative splicing. We identified protein kinase A (PKA) as the kinase that phosphorylates KDM3A serine 265 following DNA damage. This phosphorylation is important for KDM3A’s function as a scaffold joining the SWI/SNF complex and the splicing factor SRSF3 to modulate its targets. We focused on one of its targets, SAT1, and showed that regulating its alternative splicing is important for proper cell-cycle regulation following DNA damage. Our work exemplifies the importance of alternative splicing in the cellular response to the cell’s changing environment.

## RESULTS

### KDM3A regulates transcription and alternative splicing of cell-cycle genes

In order to learn how changes in the cellular environment result in chromatin structure alteration, which in turn regulates gene expression, we silenced the prominent cellular sensor histone demethylase KDM3A. Since KDM3A’s role in breast cancer progression is well established, we chose to use the breast cancer cell-line MCF7^31,32^. We began by characterizing all KDM3A-dependent expression and alternative splicing events by means of deep RNA-sequencing at a genome-wide scale using siRNA (Supplementary Fig. S1A&B). As expected of a demethylase, we detected 1075 genes that are differentially expressed when KDM3A is silenced (cut-off set at 1.5-fold); of these 3% were also alternatively spliced. Interestingly, we found a further group of 777, which did not show changes in expression, but that were alternatively spliced (Fig. 1A). This result suggests that KDM3A is a regulator of alternative splicing and that alternative splicing regulation by KDM3A, directly or indirectly, is almost exclusively independent of its role as a transcription regulator.

**Figure 1.**
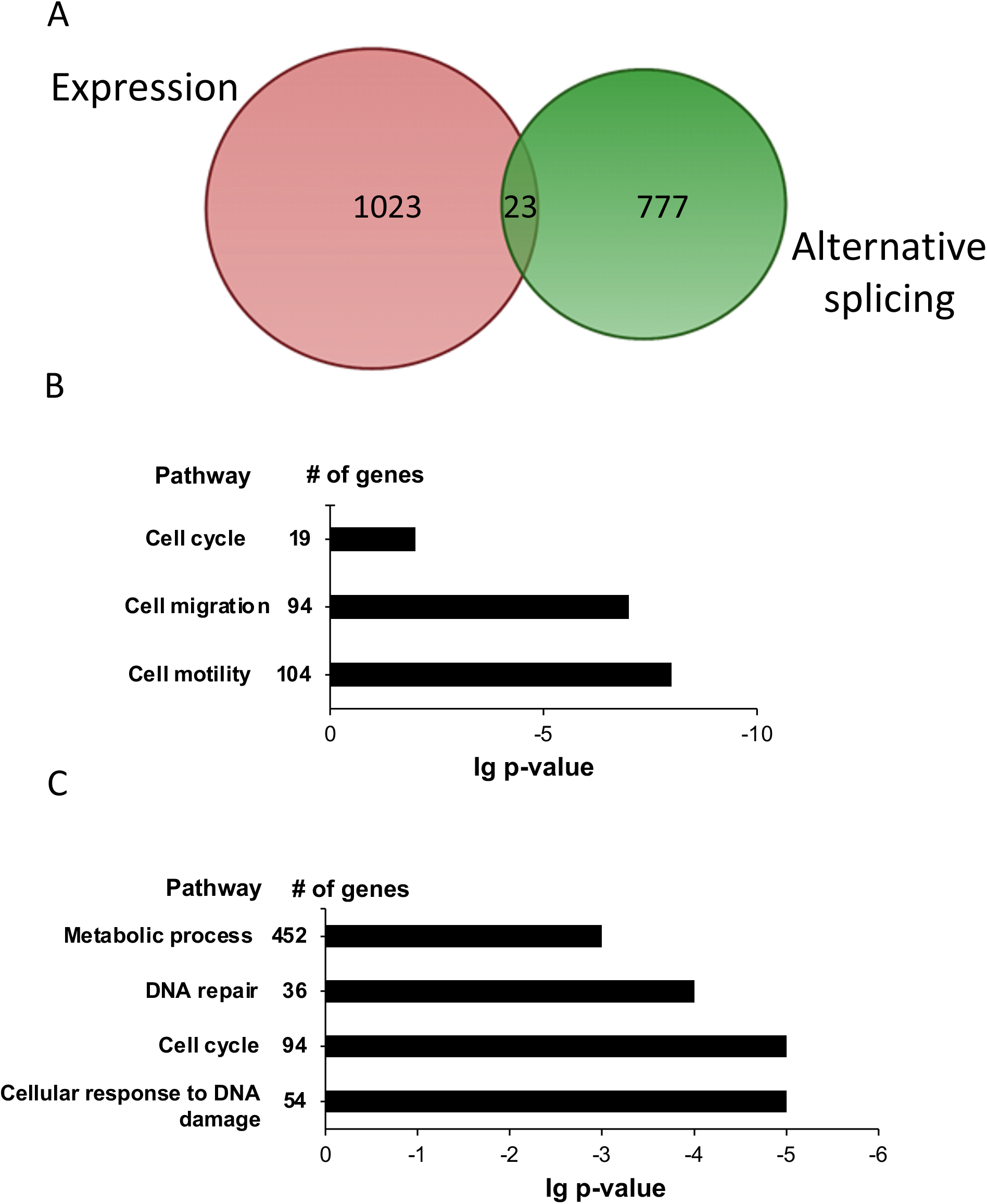
KDM3A is a regulator of transcription and alternative splicing of cell cycle and DNA damage genes. MCF7 cells were transfected with non-targeting siRNA (siNT) and siKDM3A for 72 h. RNA was extracted, libraries prepared and RNA-seq was conducted. **A**. Venn diagram representing differentially expressed genes (blue) and alternatively spliced genes (red). **B & C**. Functional analysis was conducted using the DAVID functional annotation tool (DAVID, https://david.ncifcrf.gov/) for (**B**) differentially expressed genes and (**C**) alternative splicing genes.

Among the 809 genes that were differentially spliced, we observed all types of alternative splicing events: skipped exons (SE; 573 events in 477genes), alternative 5’ splice site (A5SS; 118 events in 109 genes), alternative 3’ splice site (A3SS; 100 events in 97 genes), mutually exclusive exons (MXE; 119 events in 96 genes) and retained introns (RI; 97 events in 94 genes). We chose to validate the five top-ranking candidates (SCML, HNRNPL, PTRH2, SAT1 and ENSA) and found all to change in alternative splicing following silencing of KDM3A (Supplementary Fig. S1C-D). Using the DAVID function annotation tool (DAVID, https://david.ncifcrf.gov/)^43,44^, we found that the list of differentially expressed genes was enriched for genes with functions in cell cycle, cell migration and cell motility (Fig. 1B). The differentially alternative spliced genes were enriched for genes involved in metabolic processes, DNA repair, cell cycle, and cellular response to DNA damage (Fig. 1C). This result demonstrates that while KDM3A regulates different genes in expression and alternative splicing, the function of the genes in the two groups is similar. In addition these results suggest a new role for KDM3A following DNA damage.

The DNA damage response is a cellular system that maintains genomic stability. This system activates an enormous amount of proteins that efficiently modulate many physiological processes. The overall cellular response to DNA damage goes far beyond repair. It modulates numerous physiological processes, and alters gene expression profiles and protein synthesis, degradation and trafficking. One of its hallmarks is the activation of cell-cycle checkpoints that temporarily halt the cell-cycle while damage is assessed and repaired ^45^.

### KDM3A is phosphorylated by PKA following induction of double-strand breaks

KDM3A phosphorylation has been shown to occur following a change in temperature^17,20^. Heat stress induces KDM3A S264 phosphorylation^20^, whereas cold stress promotes KDM3A S265 phosphorylation^17^. In order to check whether KDM3A is phosphorylated following DNA damage, we began by exploring mass-spec data using phospho-site^46^. We found that KDM3A S265 is phosphorylated following treatment with etoposide, a topoisomerase II inhibitor that primarily causes double-strand DNA breaks^47^. To validate this mass-spec result, we induced double-strand DNA breaks in MCF7 cells with the radiomimetic drug neocarzinostatin (NCS). We detected KDM3A S265 phosphorylation using an antibody specific for its phospho-S265^17^. Our result shows that KDM3A S265 is phosphorylated while the total amount of KDM3A does not change following DNA damage (Fig. 2A).

**Figure 2.**
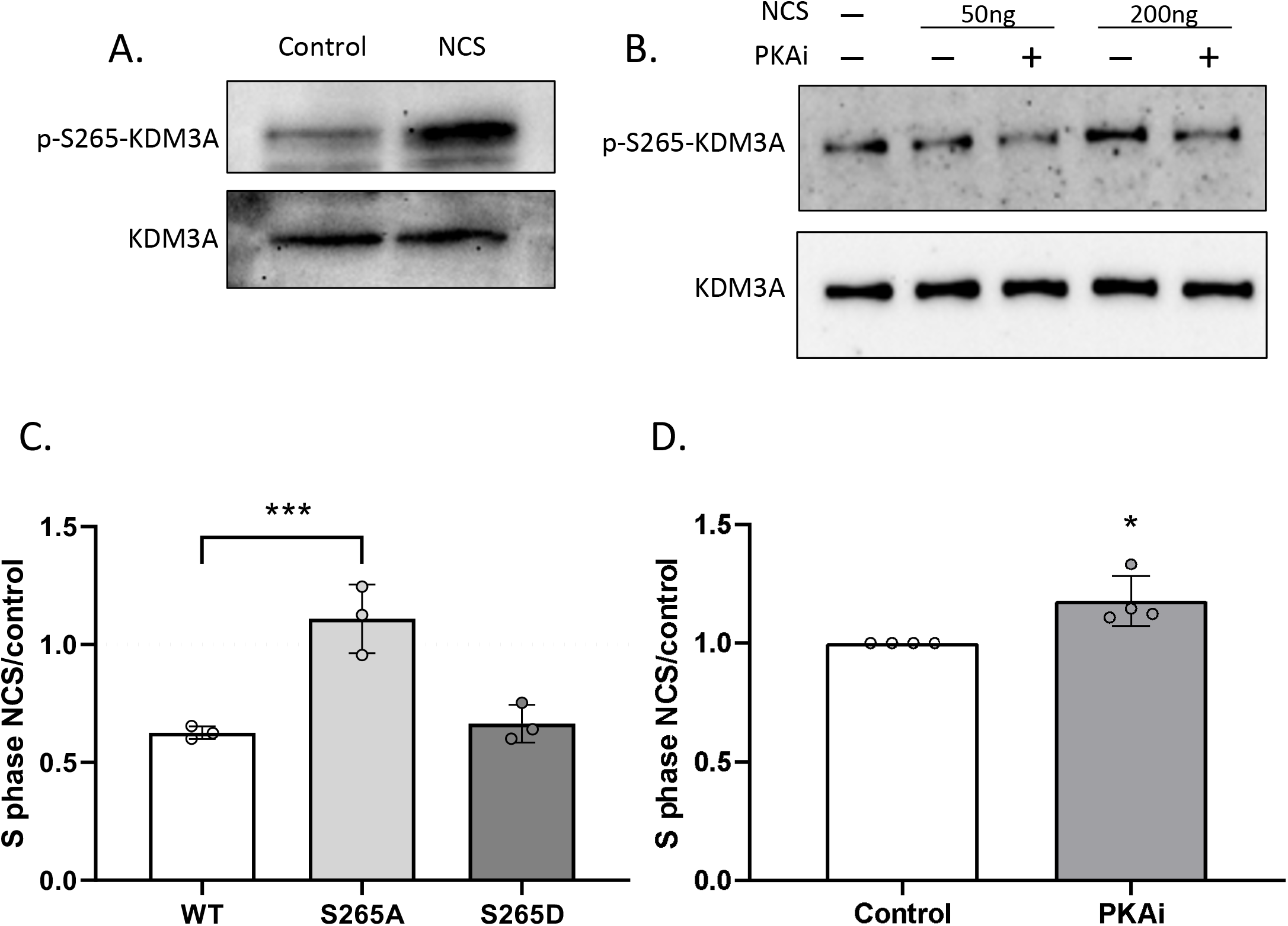
KDM3A is phosphorylated by PKA following double strand breaks. **A**. MCF7 cells treated with 200 ng/ml NCS for 1 h. Immunoblotting was conducted with the indicated antibodies. **B**. MCF7 cells were treated with PKAi for 30 min following treatment with 50 and 200 ng/ml NCS for 1 h. Immunoblotting was conducted with the indicated antibodies. **C**. MCF7 cells stably expressing either wild-type KDM3A or KDM3A mutated at serine 265 to alanine (S265A) or to aspartic acid (S265D) were incubated with 200 ng/ml NCS, and analyzed 8 h later using flow cytometry. The amount of cells accumulated in S-phase is shown as a fold-change relative to untreated cells. **D**. MCF7 cells were incubated with PKAi for 30 min followed by treatment with 200 ng/ml NCS, and analyzed 8 h later using flow cytometry. The amount of cells accumulated in S-phase is shown as a fold-change relative to untreated cells. C and D, plot represents the mean of four independent experiments and ±SD (* p<0.05; ** p<0.01).

We continued by asking which is the kinase that phosphorylates KDM3A. While the nuclear kinase ataxia telangiectasia mutated (ATM) is the main kinase activated following double-strand breaks^48^, the motif in KDM3A S265 does not match ATM’s preference (serine followed by glutamine, SQ). The motif in fact suggests that the kinase is PKA^17^. While PKA is not a bona-fide DNA damage kinase, it is known to phosphorylate ataxia telangiectasia and Rad3-related protein (ATR) following single-strand breaks^49^. To check whether PKA phosphorylates KDM3A on S265 following double-strand breaks, we incubated MCF7 cells with NCS with or without PKA inhibitor. Our results show reduced phosphorylation of KDM3A S265 following inhibition of PKA, indicating that PKA indicating that PKA, or another kinase phosphorylated by PKA, is KDM3A’s kinase (Fig. 2B).

### More cells in S phase following silencing of KDM3A or mutation at its phosphorylation site

To examine KDM3A’s role in cell cycle checkpoint following DNA damage, we silenced KDM3A using siRNA and induced double-strand breaks with NCS. We found a mild increase in percentage of cells in S-phase in cells silenced for KDM3A relative to control (Fig. S2A-C). The slight change might result from redundancy of KDM3A function with other demethylases such as KDM4C^50^.

To check if phosphorylation of KDM3A S265 is important for S-phase checkpoint, we stably expressed KDM3A with S265 mutated to alanine (S265A) or aspartic acid (S265D), mimicking the non-phosphorylated or phosphorylated form, respectively (Supplementary Fig. S2D). Comparing the cell cycle recovery in these cells to cells stably expressing WT KDM3A, we found a higher percentage of cells in S-phase in S265A but not in S265D, suggesting that KDM3A phosphorylation plays a role in cell cycle regulation (Fig. 2C and Supplementary Fig. S2E). In addition, we monitored the cell cycle of MCF7 cells whose PKA was inhibited and found a similar effect with a higher percentage of cells in S-phase (Fig. 2D and Supplementary Fig. S2F). This supported the idea that PKA and KDM3A are in the same pathway in the cellular response to DNA damage.

### SAT1, a KDM3A target in alternative splicing regulation

To study KDM3A’s role in cell cycle via alternative splicing, we chose to examine its target gene SAT1 (Supplementary Fig. S1C). SAT1 belongs to the acetyltransferase family and is a rate-limiting enzyme in the catabolic pathway of polyamine metabolism. The acetylation of spermidine and spermine by SAT1 regulates the cellular levels of free spermidine/spermine and is critical since a decrease in their concentrations inhibits cell proliferation while an excess appears to be toxic^51^. Thus the exact amount of SAT1 is important and regulated by transcription, mRNA splicing, translation, and protein stability^19^. SAT1 has seven exons, and recently it was reported that exon X (present between SAT1 exon 3 and 4) undergoes alternative splicing (Fig. 3A)^19^. Exon X codes for three stop codons and as a result the mRNA harboring exon X is a target of the nonsense-mediated mRNA decay process^19^. This is an example of an alternative splicing event that leads to a decrease in the total protein amount. Our RNA-seq followed by validation demonstrated that silencing of KDM3A promoted SAT1 exon X inclusion (Supplementary Fig. S1C and S3A-D), which decreased the amount of stable SAT1 mRNA^19^. Complementarily, overexpression of KDM3A in MCF7 cells promoted SAT1 exon X exclusion (Supplementary Fig. S3E&F), and in the same way, following NCS we observed a slight exclusion of SAT1 exon X that was abolished by inhibition of PKA (Supplementary Fig. S3G&H). This result suggests that KDM3A phosphorylation is important for SAT1 alternative splicing. To provide support for this, we monitored SAT1 exon X inclusion in our cells that stably expressed either S265A or S265D. We found that overexpression of KDM3A in its constitutively phosphorylated form promoted SAT1 exon X exclusion, similar to the alternative splicing seen following DNA damage (Supplementary Fig. I&J). Next we investigated the importance of SAT1 alternative splicing to cell cycle in MCF7. We silenced SAT1, mimicking inclusion of exon X. We found a higher percentage of cells in S-phase resembling the effect of silencing KDM3A (Fig. 3B and Supplementary Fig. S3K&L). This result suggests that KDM3A and SAT1 are working in the same pathway in the cellular response to DNA damage.

**Figure 3.**
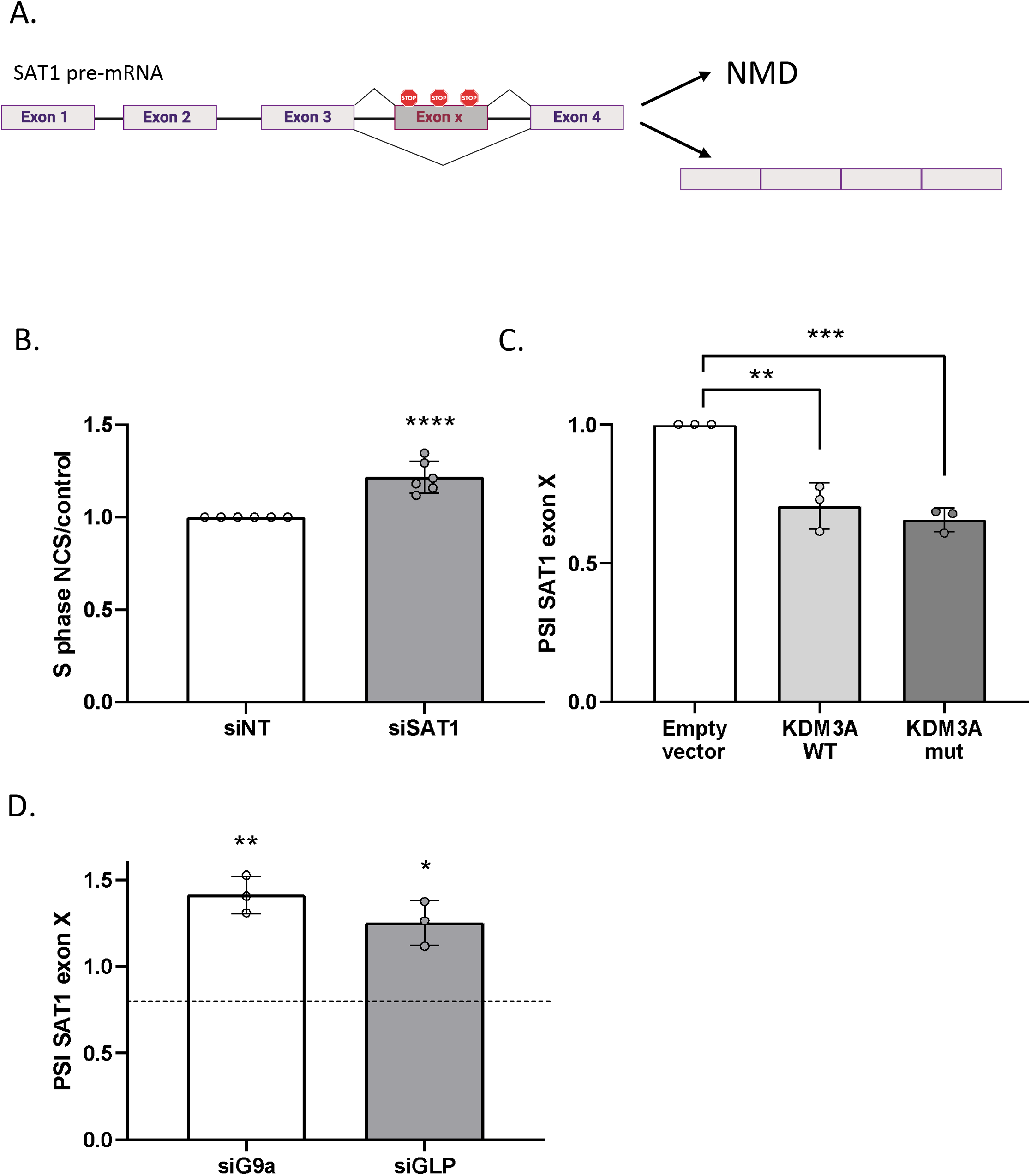
KDM3A regulates SAT1 alternative splicing independent of its demethylase enzymatic activity. **A**. Schematic representation of SAT1 alternative splicing. Rectangles: exons, black lines: introns. **B**. MCF7 cells were transfected with non-targeting siRNA (siNT) and siSAT1 for 72 h, followed by treatment with 200 ng/ml NCS, and analyzed 8 h later using flow cytometry. The amount of cells accumulated in S-phase is shown as a fold-change relative to untreated cells. **C**. MCF7 cells stably expressing either the empty vector, wild-type KDM3A or KDM3A mutated demethylase (mut). Total RNA was extracted and analyzed by real-time PCR for SAT1 exon X inclusion. PSI was calculated by SAT1 exon X relative to SAT1 total mRNA amount. **D**. MCF7 cells were transfected with non-targeting siRNA (siNT) and siG9a or siGLP for 72 h. Total RNA was extracted and analyzed by real-time PCR for SAT1 exon X inclusion. PSI was calculated by SAT1 exon X relative to SAT1 total mRNA amount. Values represent averages of three independent experiments ±SD (* p<0.05; ** p<0.01; *** p<0.001; **** p<0.0001).

### KDM3A serves as a scaffold to regulate SAT1 alternative splicing

To understand how KDM3A regulates SAT1 alternative splicing, we first asked whether the demethylase activity of KDM3A is critical for its regulation of alternative splicing. To this end, we stably expressed KDM3A with a mutated demethylase domain and silenced endogenous KDM3A in MCF7 cells. We found that overexpression of either WT or the demethylase mutant KDM3A reduced SAT1 exon X inclusion (Fig. 3C and Supplementary Fig. S3M&N). This result indicates that KDM3A’s demethylase activity is not essential for alternative splicing regulation and suggests that KDM3A serves as a scaffold to regulate SAT1 alternative splicing. We then silenced the H3K9 methyltransferases G9a and GLP and the H3K9 methylations readers: HP1α, HP1β and HP1γ and monitored SAT1 alternative splicing. We observed SAT1 exon X inclusion when G9a or GLP were silenced (Fig. 3D and Supplementary Fig. 3O), similar to the results seen when silencing KDM3A. No change in SAT1 alternative splicing was observed following silencing of H3K9 methylation readers (Supplementary Fig. P&Q). This result suggests that the methylation state of H3K9 might recruit KDM3A, but that KDM3A does not serve as an eraser of the modification but rather as a scaffold for splicing factor binding.

### SRSF3 and hnRNPF promote exclusion of SAT1 exon X

We hypothesize that KDM3A serves as a scaffold to recruit a splicing factor to exclude SAT1 exon X. Such splicing factors could be those known to act as an adaptor system connecting DNA and histone methylation to the spliceosome SRSF1, SRSF3, SRSF6 and SF3B1^42^ and hnRNPF, which was shown to bind KDM3A and regulate alternative splicing of AR variant 7 (AR-V7)^28^. We silenced each of these splicing factors and found that both SRSF3 and hnRNPF promote exclusion of SAT1 exon X similar to KDM3A (Fig. 4A and Supplementary Fig. S4A-C). This suggests that SRSF3 and hnRNPF might be working in the same pathway as KDM3A in the regulation of SAT1 alternative splicing.

**Figure 4.**
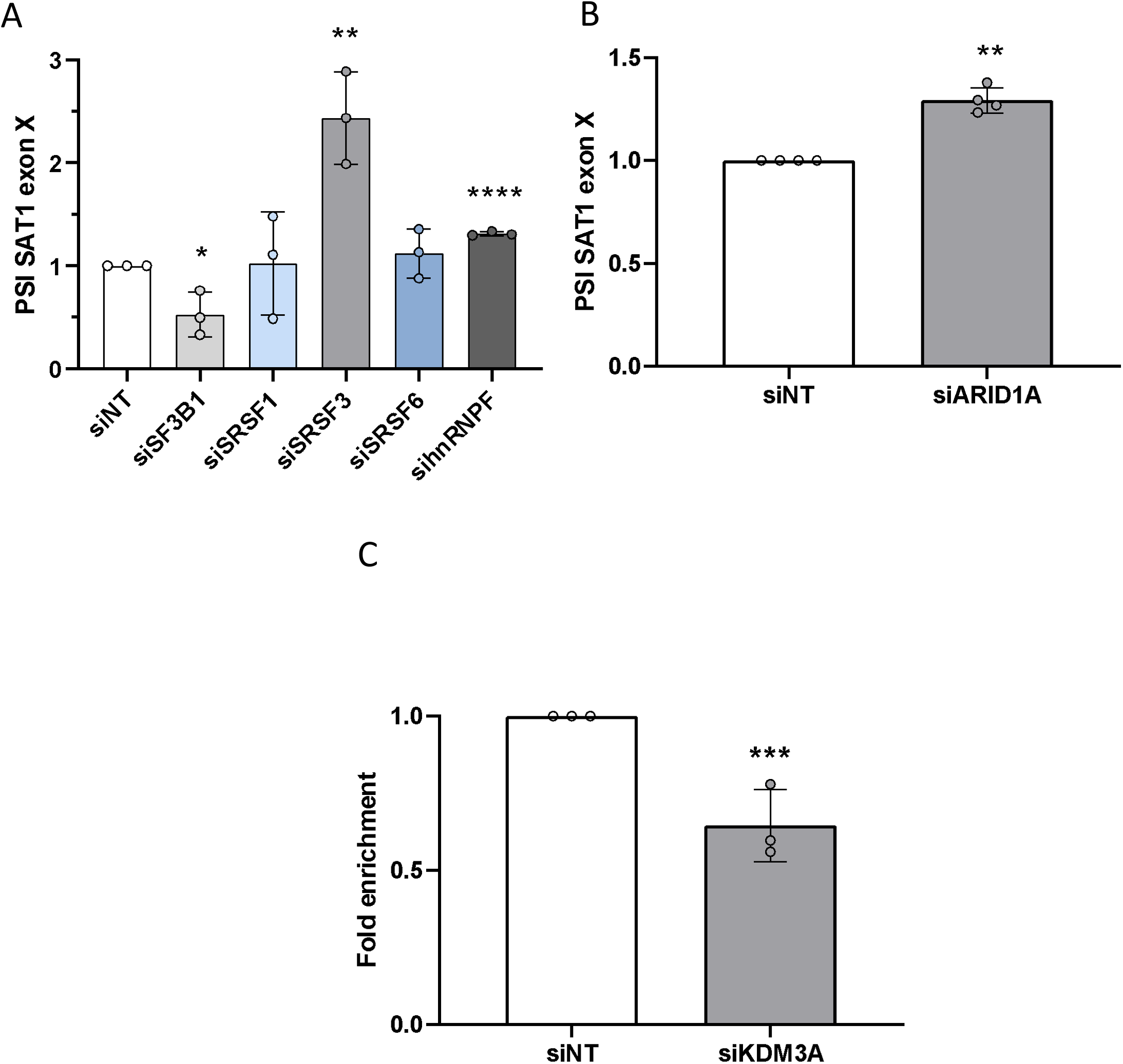
KDM3A allows for SRSF3 binding to SAT1 pre-mRNA. **A**. MCF7 cells were transfected with non-targeting siRNA (siNT) or siSF3B1, siSRSF1/3/6 and sihnRNPF for 72 h. Total RNA was extracted and analyzed by real-time PCR for SAT1 exon X inclusion. PSI was calculated by SAT1 exon X relative to SAT1 total mRNA amount. **B**. MCF7 cells were transfected with non-targeting siRNA (siNT) or siARID1A for 72 h. Total RNA was extracted and analyzed by real-time PCR for SAT1 exon X inclusion. PSI was calculated by SAT1 exon X relative to SAT1 total mRNA amount. **C**. MCF7 cells stably expressing FLAG-KDM3A-V5 were treated with 200ng/ml NCS for half an hour. KDM3A was immunoprecipitated using the Flag tag, and detected with V5 (for KDM3A) and SRSF3. * long exposure. **D**. RNA-IP of SRSF3 in MCF7 cells that were transfected with non-targeting siRNA (siNT) or siKDM3A for 72 h. Real-time PCR analysis of SAT1 intron 3 (near exon X) relative to input. Values represent averages of three independent experiments ±SD (* p<0.05; ** p<0.01; **** p<0.0001).

### The SWI/SNF subunit ARID1A promotes SAT1 exon X exclusion

KDM3A phosphorylation on S265 following β-adrenergic stimulation regulates gene expression via the SWI/SNF complex^17^. The SWI/SNF subunit Brm was previously shown to promote exon inclusion of several genes^52^. This led us to ask whether the SWI/SNF complex has a role in SAT1 alternative splicing. To check our hypothesis, we asked whether ARID1A, a SWI/SNF subunit binding to phospho-KDM3A^17^, has a role in SAT1 alternative splicing. To this end, we silenced ARID1A and found that it promoted SAT1 exon X exclusion (Fig. 4B and Supplementary Fig. S4D-F). Since ARID1A, KDM3A, and SRSF3 promote SAT1 exon X exclusion, it is possible that they cooperate in the alternative splicing of SAT1 exon X, presumably by forming an adaptor system connecting chromatin to the spliceosome.

### KDM3A is important for SRSF3 binding to SAT1 pre-mRNA

To check if KDM3A is needed for SRSF3 binding to SAT1 pre-mRNA, we performed RNA-IP for SRSF3 following silencing of KDM3A in MCF7 cells (Supplementary Fig. S4G&H). Monitoring SAT1 pre-mRNA relative to input, we found silencing of KDM3A led to a reduction of approximately 75% in SRSF3 binding to SAT1 pre-mRNA (Fig. 4C). This result supports the hypothesis that the two proteins functionally interact in the regulation of SAT1’s alternative splicing.

## MATERIALS AND METHODS

### Cell culture

MCF7 (ATCC Number: HTB-22) and HEK293T (ATCC Number: CRL-3216) cells were grown in Dulbecco’s modified Eagle’s medium (DMEM) supplemented with 10% fetal bovine serum; cell-lines were maintained at 37°C and 5% CO^2^ atmosphere.

### RNA interference

A pool of four siRNA oligomers per gene against KDM3A, SRSF1, SRSF3, SRSF6, HP1α, HP1β, HP1γ and SF3B1 were purchased from Dharmacon; SAT1, HNRNPF and ARID1A esiRNA were purchased from Sigma. MCF7 cells were grown to 20–30% confluence and transfected with siRNA using TransIT-X2 transfection reagent following the manufacturer’s instructions. After 24 h of incubation, cell culture media was refreshed and then incubated for an additional 48-72 h. Knockdown efficiencies were verified by qPCR.

### qRT-PCR

RNA was isolated from cells using the GENEzol TriRNA Pure Kit (GeneAid). cDNA synthesis was carried out with the Quanta cDNA Reverse Transcription Kit (QuantaBio). Then, qPCR was performed with the iTaq Supermix (BioRad) on the Biorad iCycler. The comparative Ct method was employed to quantify transcripts, and delta Ct was measured in triplicate. Primers used to amplify the target genes are provided in Supplementary Table 1.

### RNA-seq

RNA ScreenTape kit (catalog #5067-5576; Agilent Technologies), D1000 ScreenTape kit (catalog #5067-5582; Agilent Technologies), Qubit RNA HS Assay kit (catalog # Q32852; Invitrogen), and Qubit DNA HS Assay kit (catalog #32854; Invitrogen) were used for each specific step for quality control and quantity determination of RNA and library preparation. For mRNA library preparation: TruSeq RNA Library Prep Kit v2 was used (Illumina). In brief, 1 µg was used for the library construction; library was eluted in 20 µL of elution buffer. Libraries were adjusted to 10 mM, and then 10 µL (50%) from each sample was collected and pooled in one tube. Multiplex sample pool (1.5 pM including PhiX 1.5%) was loaded in NextSeq 500/550 High Output v2 kit (75 cycles cartridge and 150 cycles cartridge; Illumina). Run conditions were in paired end (43 × 43 bp and 80 × 80 bp, respectively) and loaded on NextSeq 500 System machine (Illumina). We used rMATS (version3.2.5)^53^ to identify differential alternative splicing events between the two sample groups corresponding to all five basic types of alternative splicing patterns. Briefly, rMATS uses a modified version of the generalized linear mixed model to detect differential alternative splicing from RNA-seq data with replicates, while controlling for changes in overall gene expression levels. It accounts for exon-specific sequencing coverage in individual samples as well as variation in exon splicing levels among replicates. For each alternative splicing event, we used the calculation on both the reads mapped to the splice junctions and the reads mapped to the exon body. The results of rMATS analysis gave low numbers of significant AS events with p-value 0.05 and a cut off at FDR < 5%. Thus, one replicate was excluded from the analysis, and then specific parameters were identified manually; with inclusion level difference between groups = 2, and average count >10.

### Viral infection

Retroviral or lentiviral particles were produced by expressing KDM3A-P S265 WT or mutants (S265A, S265D)^17^ or p-Lenti-V5-KDM3A-WT (addgene #82331) or demethylase MUT (addgene #82332) plasmids respectively in the HEK293T packaging cell-line using TransIT-X2 transfection reagent. MCF7 cells were infected with viral particles and stable integrations were selected.

### Immunoblotting and immunoprecipitation

For immunoblotting cells were harvested and lysed with RIPA lysis buffer, and the extracts were run on a 4–12% Bis-Tris gel and transferred onto a polyvinylidene difluoride membrane. For immunoprecipitation cells were washed twice with ice-cold PBS, harvested and lysed for 30 min on ice in 0.5% NP40, 150 mM NaCl, 50 Mm Tris pH7.5, and 2 mM MgCl2 supplemented with protease, phosphatase and RNase inhibitors. Supernatants were collected after centrifugation at 21, 000 g for 20 min. Supernatants were immunoprecipitated for 2 h with 40 ul of anti-Flag M2 Magnetic Beads (M8823, Sigma Aldrich). Beads were washed sequentially for 5 min. Beads were boiled in sample buffer and loaded onto the gel for analysis. The samples were subjected to standard immunoblotting analysis using polyvinylidene difluoride membranes and enhanced chemiluminescence.

### Antibodies

Antibodies used for immunoblotting were KDM3A-Ser265 (Kindly supplied by Dr. Juro Sakai, RCAST, University of Tokyo), KDM3A (cat# ab106456, Abcam), Arid1A (cat# sc-32761, Santa Cruz Biotechnology), SRSF3 (cat# sc-13510, Santa Cruz Biotechnology), SAT1 (D1T7M, cat# 61586, Cell Signaling Technology).

### Flow cytometry

Cells were trypsinized, washed with PBS, fixed overnight at -20°C with 70% ethanol in PBS, washed with PBS, and left for 30 min at 4°C. The cell suspension was then incubated with PBS containing 5 µg/ml DNase-free RNase and stained with propidium iodide (PI). Data was acquired using LSRII Fortessa analyser machine at 10,000 events/sample. The percentage of cells in each cell cycle phase was determined using Flow Jo software.

### Chemical reagents

Neocarzinostatin (NCS) was obtained from Sigma (Ca# 067M4060V). PKA inhibitor was obtained from Sigma (P6062).

### RNA-immunoprecipitation (RNA-IP)

Cells were washed with ice-cold PBS, harvested, and lysed for 30 min on ice in a buffer containing 1% NP40 150 mM NaCl, 50 mM Tris pH = 7.5, and 1 mM EDTA supplemented with protease and RNAsin inhibitors followed by sonication in an ultrasonic bath (Qsonica, Q800R2 Sonicator) for 6 cycles of 5 s ON and 30 s OFF. Supernatants were collected after centrifugation at 21,000 g for 20 min. Antibody was added for 2 h at 4°C. Protein A and G sepharose beads were added for an additional 1 h. Beads were washed 4 times and GeneZol added for RNA extraction. Serial dilutions of the 10% input cDNA (1:5, 1:25, 1:125) were analyzed by SYBR-Green real-time qPCR. The oligonucleotide sequences used are listed in Supplementary Table S1.

## DISCUSSION

Chromatin structure is constantly changing in response to the cell’s environment. These dynamic alterations are the basis of a changing cellular transcriptome that in turn adjust the cell to the new environment. Here we suggest a transcriptome change by alternative splicing giving rise to isoforms that are better able to respond to the environment. To this end we studied the role of the signal-sensing scaffold KDM3A. While KDM3A’s role in temperature sensing is well established, we have now found that it is phosphorylated following DNA damage to regulate alternative splicing, as well as the expression of cell cycle genes.

DNA damage has been shown multiple times by us and others to regulate the transcriptome and specifically alternative splicing^54-57^. Here we add that the mediator of the alternative splicing regulation can be chromatin proteins that are a known part of the massive cellular response to DNA damage. Splicing factors were also demonstrated to be part of the signal transduction following DNA damage. In particular SRSF3 was demonstrated to serve as a regulator of genome integrity and cell cycle^58-61^. Our work demonstrates that chromatin alternations following DNA damage could serve as a scaffold to recruit SRSF3 to specific cell cycle genes. This observation could add a level of regulation to splicing factors’ preference for their targets following a cellular environment change.

Histone modifications over alternatively spliced gene regions have previously been shown to act as recognition sites for epigenetic adaptor proteins, which in turn recruit splicing factors^4,33^. Here we demonstrate that phosphorylated S265 KDM3A itself could serves as a scaffold for splicing factor recruitment with the SWI/SNF subunit ARID1A. Another subunit of the SWI/SNF complex, Brm, was indicated to regulate alternative splicing by promoting inclusion of exons via slowing down of RNA polymerase II elongation^52^. However, our results show that ARID1A promotes exclusion of SAT1 exon X. This suggest that the mechanism of the SWI/SNF complex on the SAT1 gene is different to that described before.

Our results demonstrate that the H3K9me2 methyltransferases, G9a and GLP, and its demethylase, KDM3A, both promote exclusion of SAT1 exon X. This result is of special interest and provide strong evidence that KDM3A serves as the scaffold, and not the H3K9 modification, for recruiting SRSF3. Our previous work found HP1γ to bind H3K9me2 and recruit the splicing factor SRSF1^33^. This model suggests that KDM3A enzymatic activity will abolish this adaptor system and will promote the opposite alternative splicing outcome. However, SAT1 is a proof that G9a and KDM3A can work in a similar way to regulate alternative splicing. A deeper investigation is needed to learn the fine tuning of each of these chromatin factors.

Demethylation of H3K9 by KDM3A was shown to promote alternative splicing of AR7 by recruiting hnRNPF^28^. This work describes a mechanism that is distinct from the one we describe here as it requires KDM3A’s enzymatic activity and our work shows that the methylase mutant of KDM3A can also serve as an alternative splicing regulator. Our results shed light on the many mechanisms by which chromatin can regulate alternative splicing.

## DATA AVAILABILITY

The RNA-seq data has been deposited to the Gene Expression Omnibus (GEO) with the dataset identifier GSE141948.

## ACKNOWLEDGEMENTS

We thank Prof. Juro Sakai and Prof. Takeshi Inagaki for plasmids and p-KDM3A antibody. We thank Abed Nasereddin and Idit Shiff from the Genomic Applications Laboratory, The Core Research Facility, The Faculty of Medicine - Ein Kerem, The Hebrew University of Jerusalem, Israel, for their professional advice and RNA-seq service.

This work was in part supported by the Alon Award by the Israeli Planning and Budgeting Committee (PBC) and the Israel Science Foundation (ISF 1154/17). The RNA-seq reagents were funded by a Core Lab Grant Program launched by Danyel Biotech, the authorized Illumina Channel Partner in Israel.

## Figure Legends

**Supplementary Figure S1.**
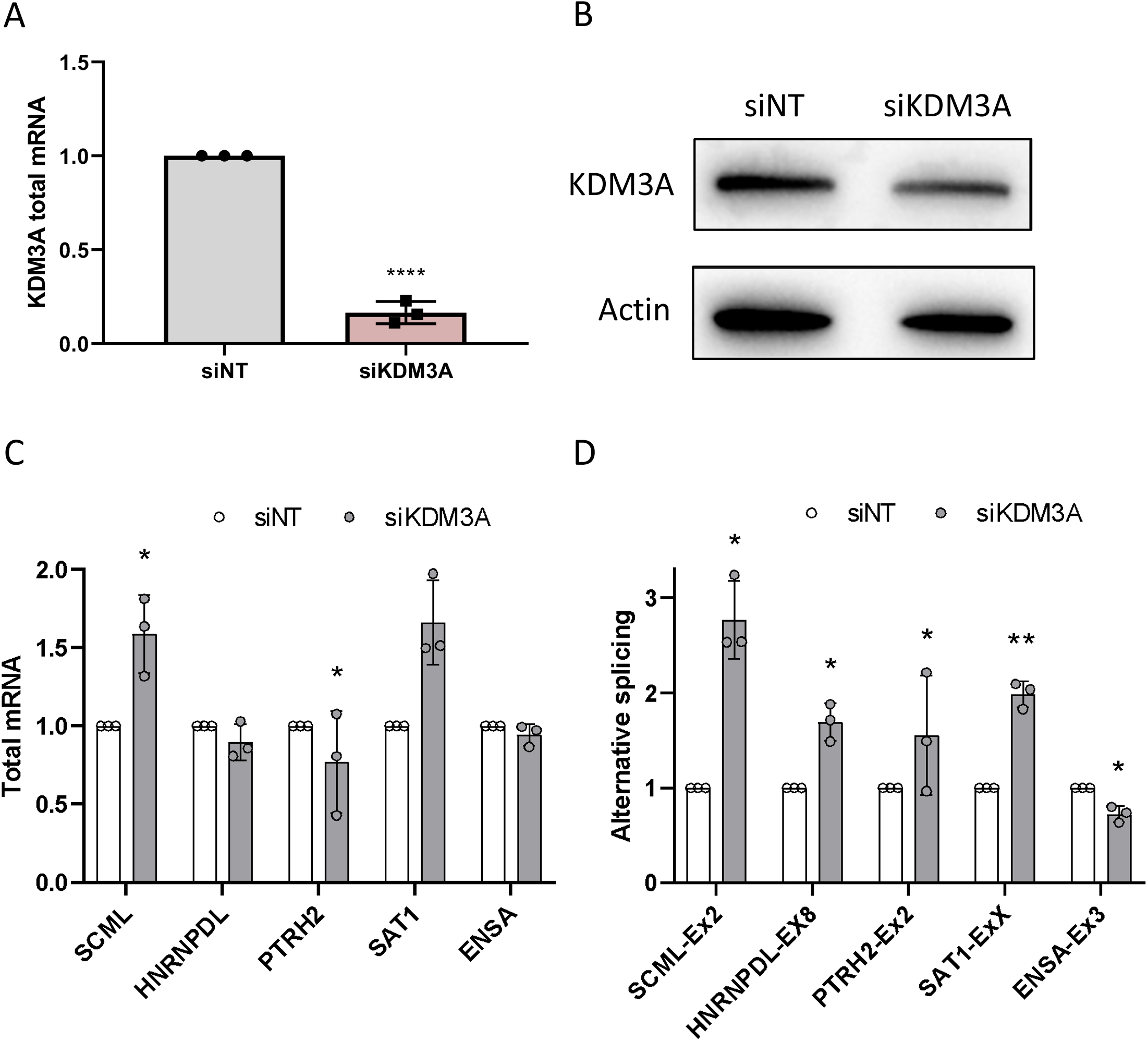
**A-D**: MCF7 cells were transfected with non-targeting siRNA (siNT) and siKDM3A for 72 h. Total RNA was extracted and analyzed by real-time PCR for total mRNA amount of KDM3A (**A**). Immunoblotting was conducting using the indicated antibodies (**B**). Five genes (HNRNPDL, SAT1, SCML, PTRH2 and ENSA) were chosen for validation and total RNA was extracted and analyzed by real-time PCR for their total mRNA amount relative to *CycloA* reference gene (**C**); PSI of indicated genes was analyzed by real-time PCR relative to the total amount of that gene (**D**). Values represent averages of three independent experiments ±SD (* p<0.05; ** p<0.01; **** p<0.0001; Student’s t test comparing to siNT).

**Supplementary Figure S2.**
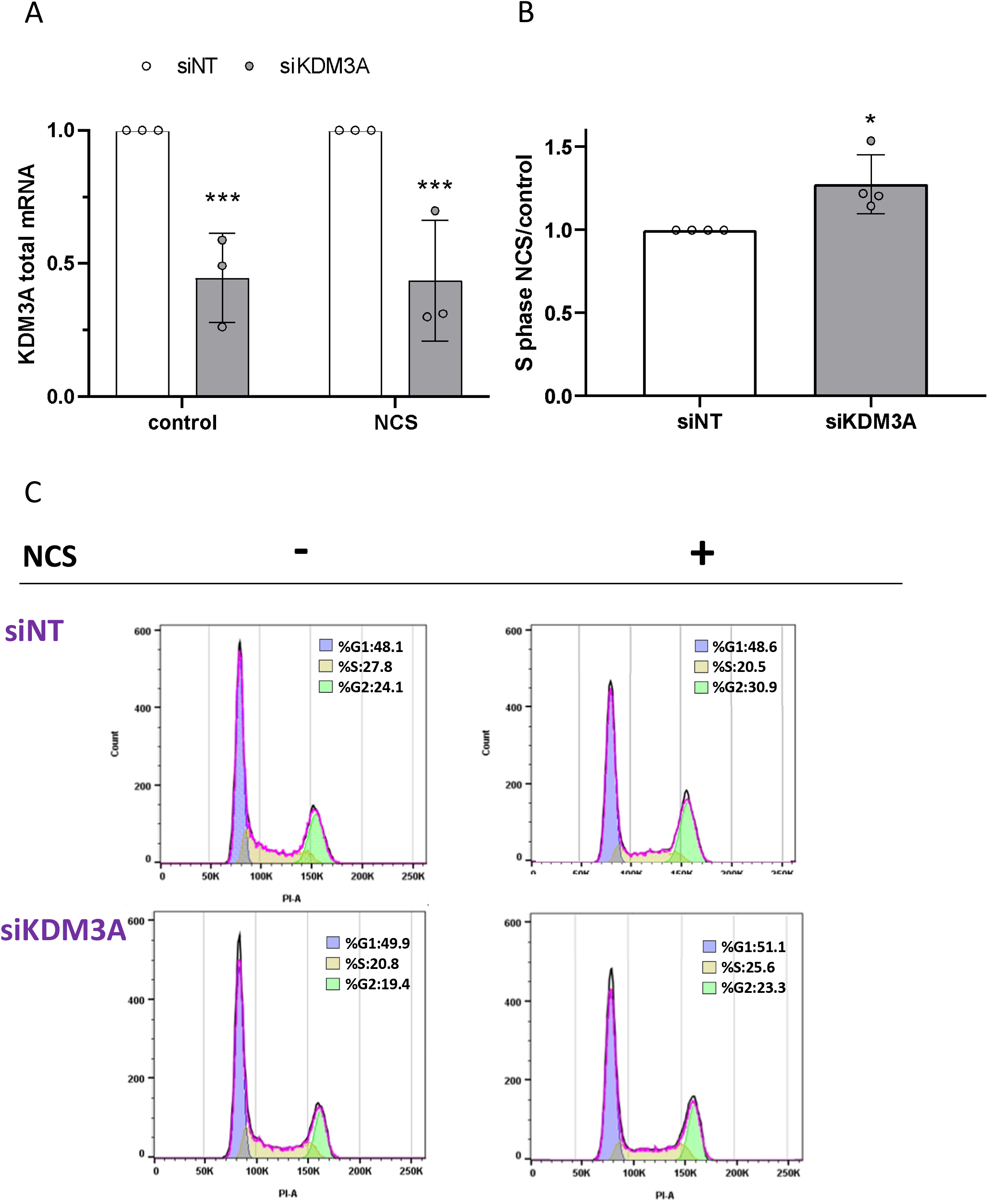
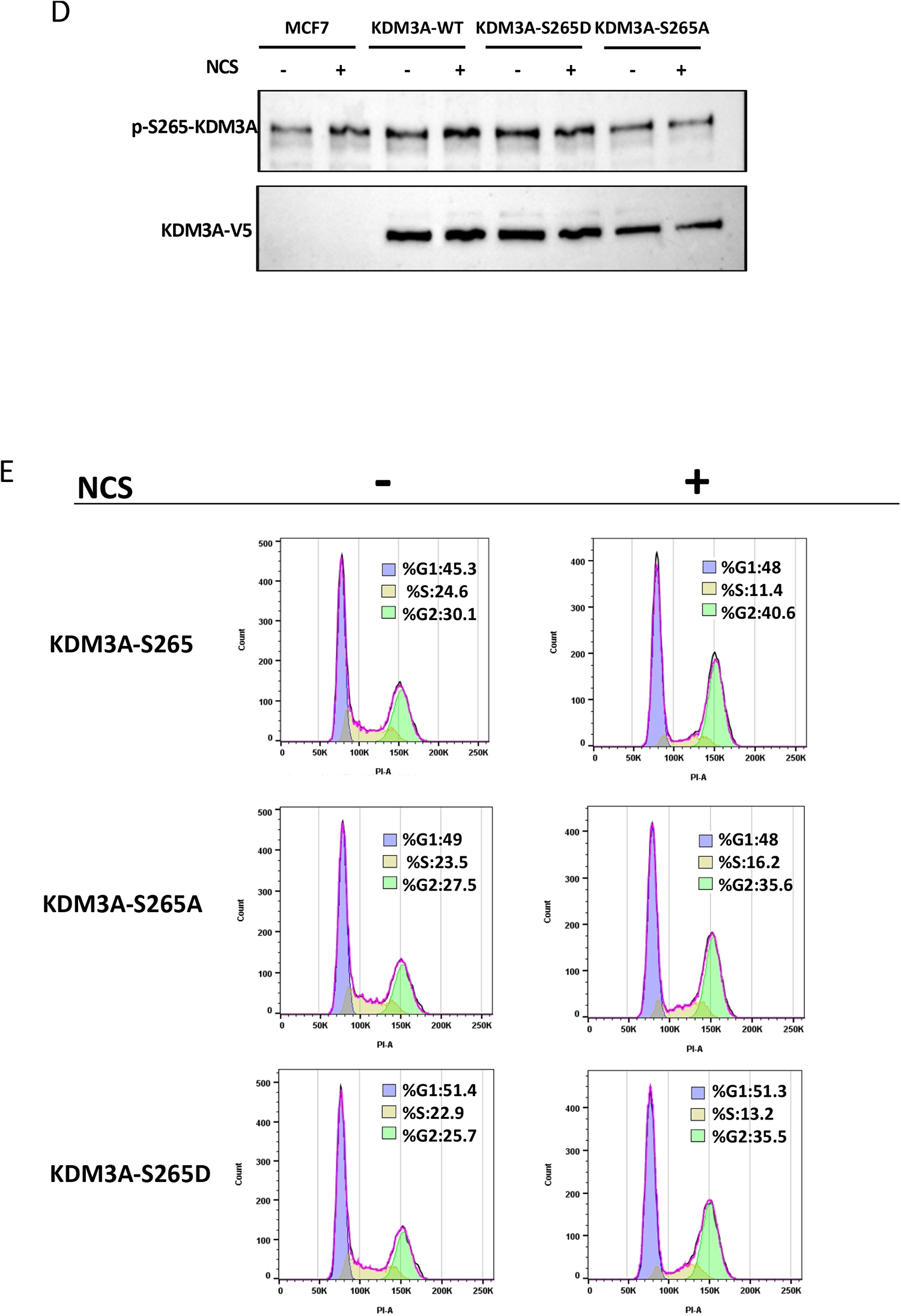
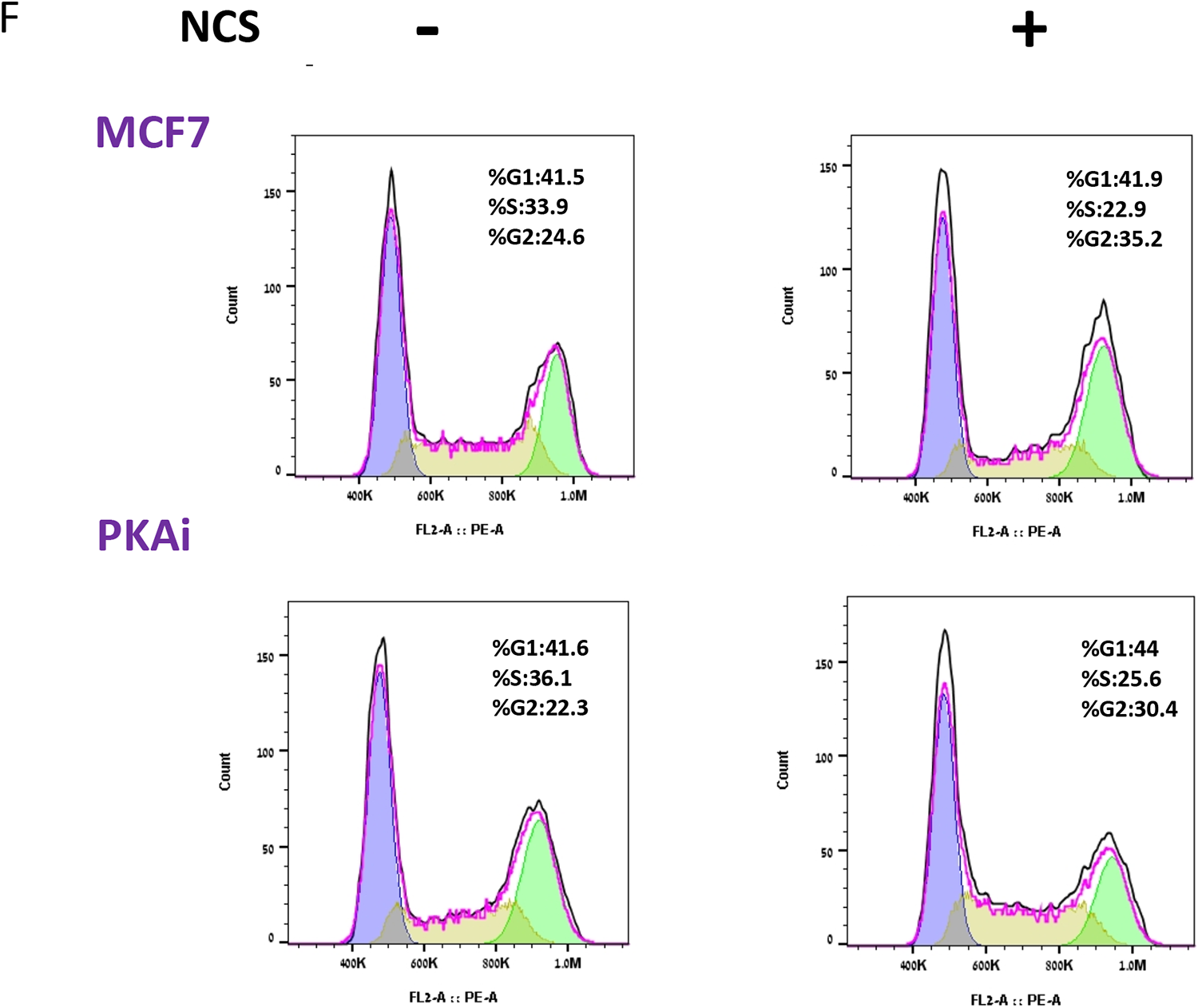
**A-D**. MCF7 cells were transfected with non-targeting siRNA (siNT) and siKDM3A for 72 h and treated with 200 ng/ml NCS for 8 h. Total RNA was extracted and analyzed by real-time PCR for total mRNA amount of KDM3A relative to *CycloA* reference gene (**A**). Cells were analyzed using flow cytometry. The amount of cells accumulated in S-phase is shown as a fold-change relative to untreated cells (**B**). Values in **A** & **B** represent averages of three independent experiments ±SD (* p<0.05; *** p<0.005; Student’s t test comparing to siNT). A population histogram for one representative experiment is shown (**C**). **D&E**. Protein was collected from MCF7 cells stably expressing either wild-type KDM3A or KDM3A mutated at serine 265 to alanine (S265A) or to aspartic acid (S265D). Immunoblotting was conducted with the indicated antibodies (**E**) and cells were treated with 200 ng/ml NCS for 8 h and analyzed using flow cytometry. A population histogram for one representative experiment is shown (**F**).

**Supplementary Figure S3:**
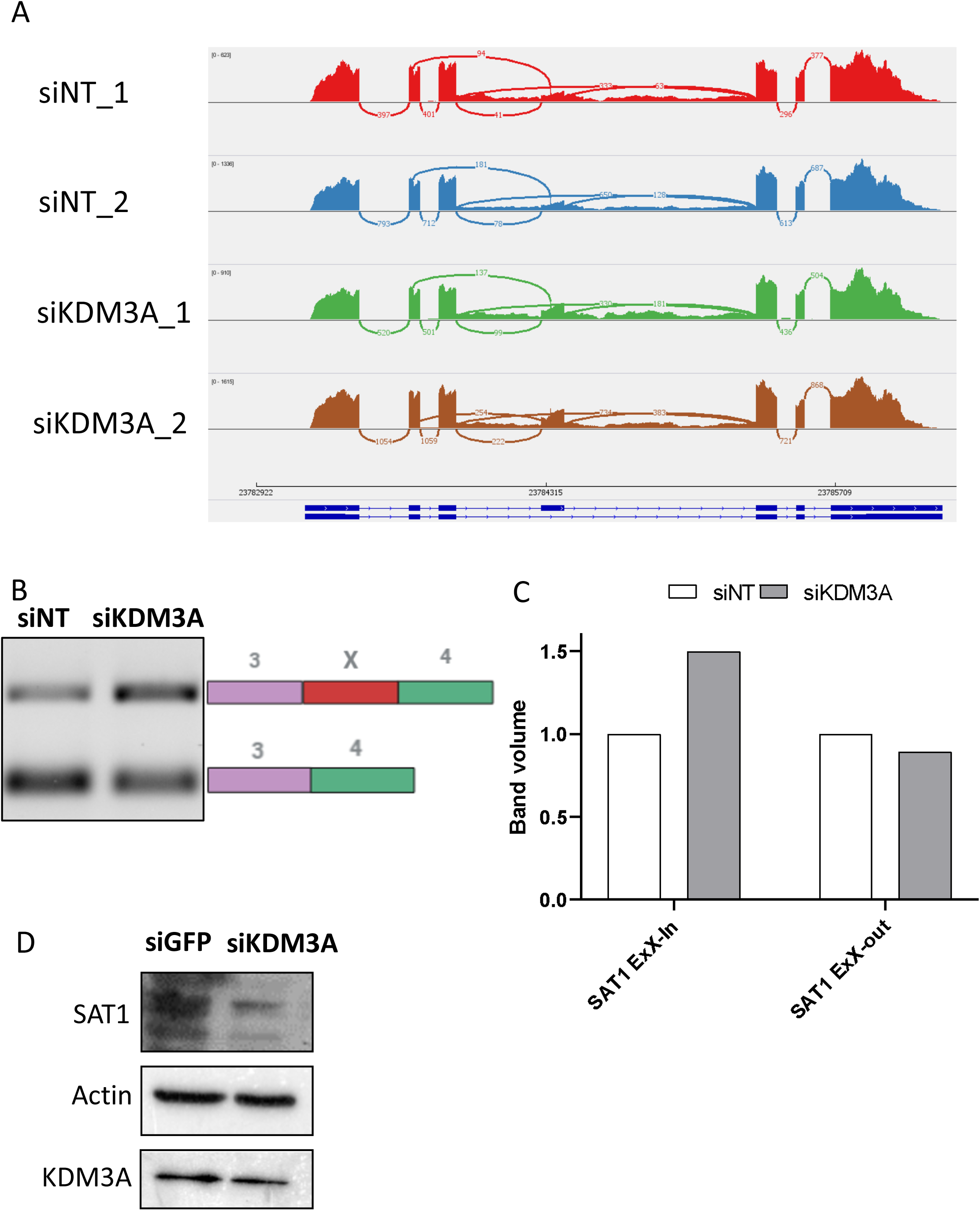
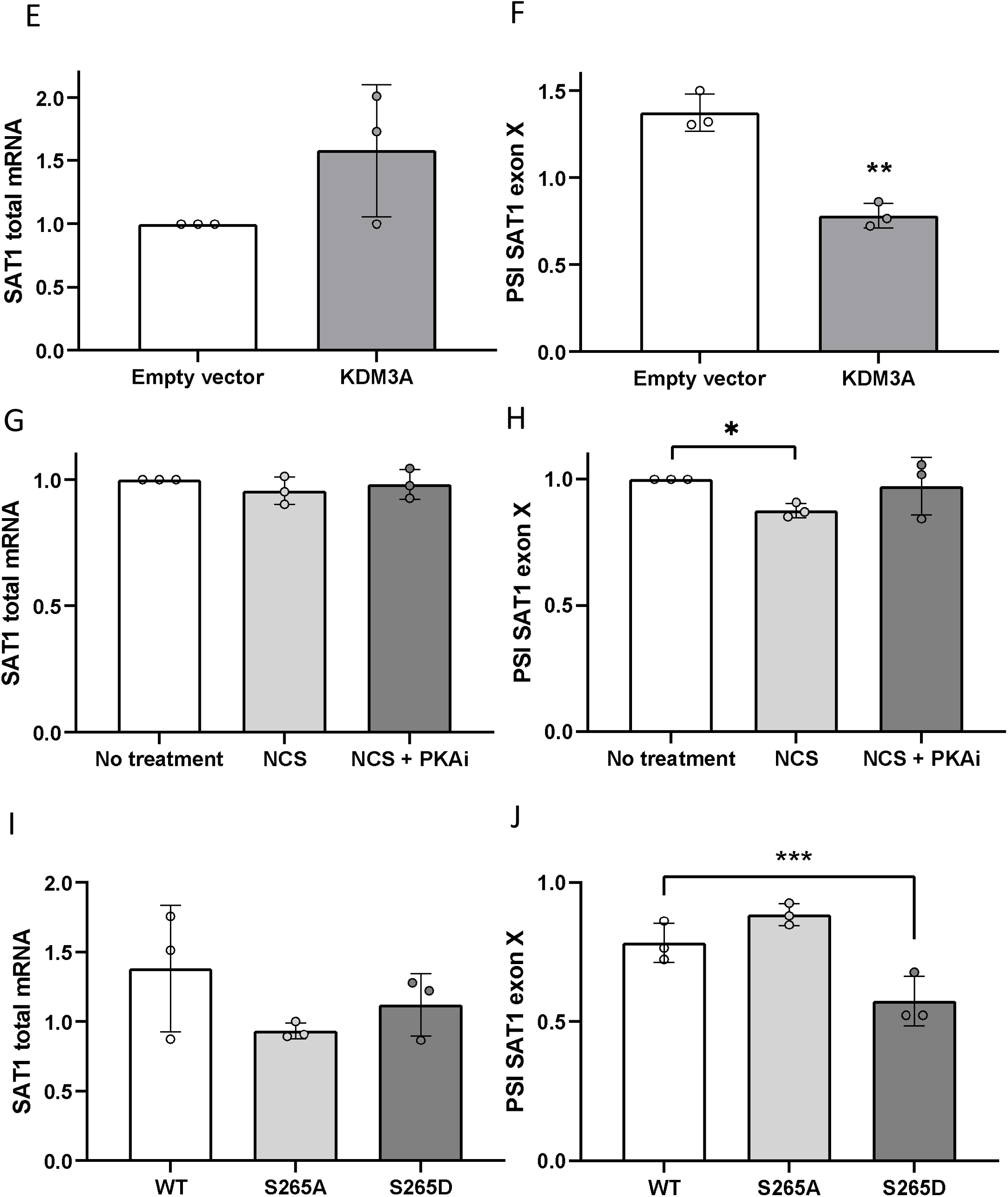
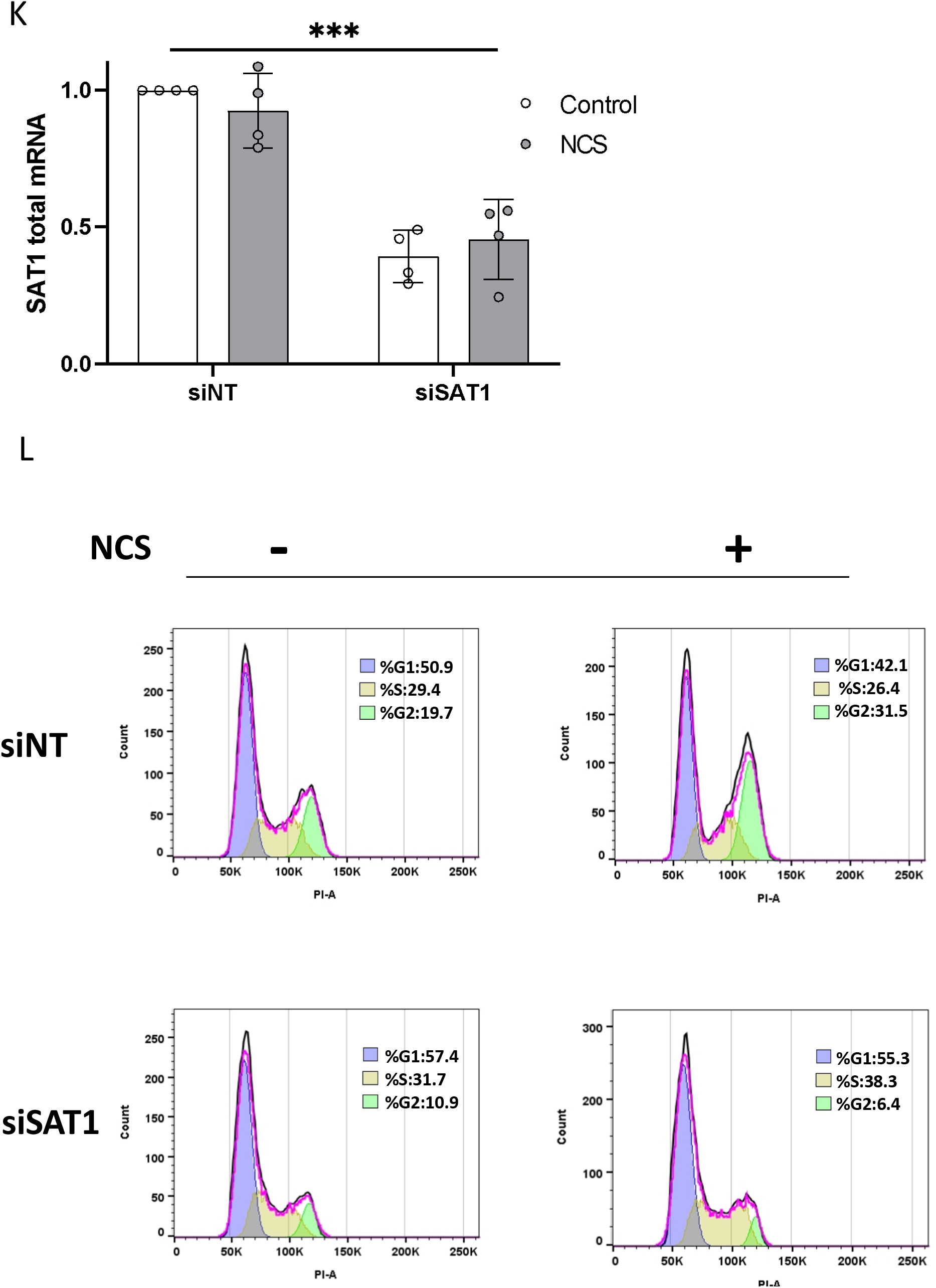
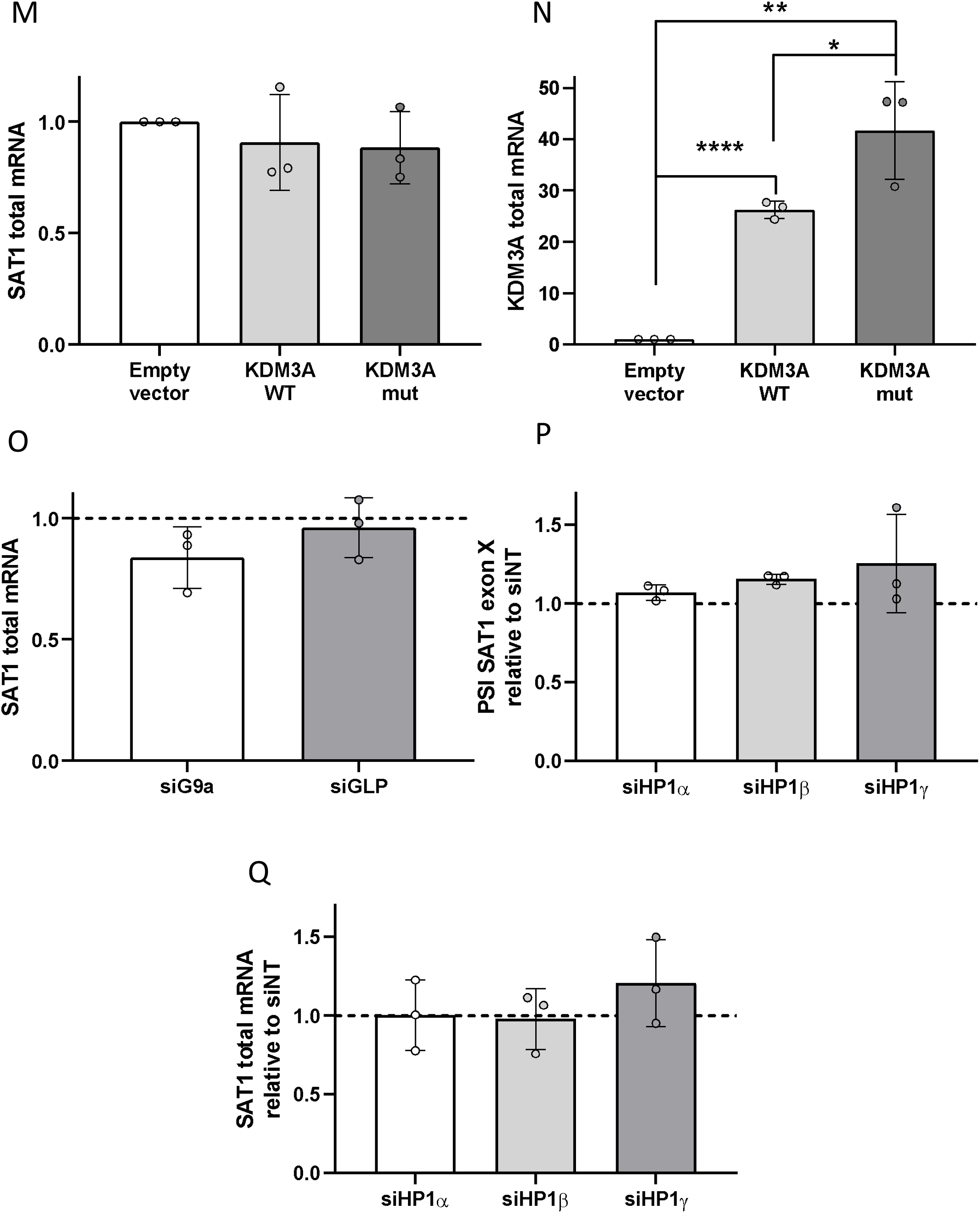
**A**. RPKM for each alignment track are shown as sashimi plots (Katz *et al*. 2015). Arcs denote splice junctions, quantified in spanning reads, as specified near each exon junction of *SAT1* gene. **B & C**. SAT1 alternative splicing in MCF7 cells stably expressing either the empty vector or wild-type KDM3A. Total RNA was extracted and analyzed by real-time PCR for KDM3A relative to *CycloA* reference gene (**B**) and for SAT1 exon X inclusion. PSI was calculated by SAT1 exon X relative to SAT1 total mRNA amount (**C**). **D & E**. MCF7 cells were treated with PKAi for 30 minutes following by treatment with 200 ng/ml NCS for 1 h. Total RNA was extracted and analyzed by real-time PCR for SAT1 relative to *CycloA* reference gene (**D**) and for SAT1 exon X inclusion. PSI was calculated by SAT1 exon X relative to SAT1 total mRNA amount (**E**). **F & G**. SAT1 alternative splicing in MCF7 cells stably expressing either wild-type KDM3A or KDM3A mutated at serine 265 to alanine (S265A) or to aspartic acid (S265D). Total RNA was extracted and analyzed by real-time PCR for SAT1 relative to *CycloA* reference gene (**F**) and for SAT1 exon X inclusion. PSI was calculated by SAT1 exon X relative to SAT1 total mRNA amount (**G**). **H & I**. MCF7 cells were transfected with non-targeting siRNA (siNT) and siSAT1 for 72 h, total RNA was extracted and analyzed by real-time PCR for SAT1 relative to *CycloA* reference gene (**H**). Population histogram for one experiment is presented (**I**). **J & K**. SAT1 and KDM3A expression in MCF7 cells stably expressing either the empty vector, wild-type KDM3A or KDM3A mutated demethylase (mut). Total RNA was extracted and analyzed by real-time PCR for SAT1 relative to *CycloA* reference gene (**J**) and real-time PCR for KDM3A relative to *CycloA* reference gene (**K**). **L**. MCF7 cells were transfected with non-targeting siRNA (siNT) or siG9a, siGLP for 72 h. Total RNA was extracted and analyzed by real-time PCR for SAT1 relative to *CycloA* reference gene. **M & N**. MCF7 cells were transfected with non-targeting siRNA (siNT) or siHP1α, β, γ for 72 h, total RNA was extracted and analyzed by real-time PCR for SAT1 relative to *CycloA* reference gene (**M**) and for SAT1 exon X inclusion. PSI was calculated by SAT1 exon X relative to SAT1 total mRNA amount (**N**).

**Supplementary Figure S4:**
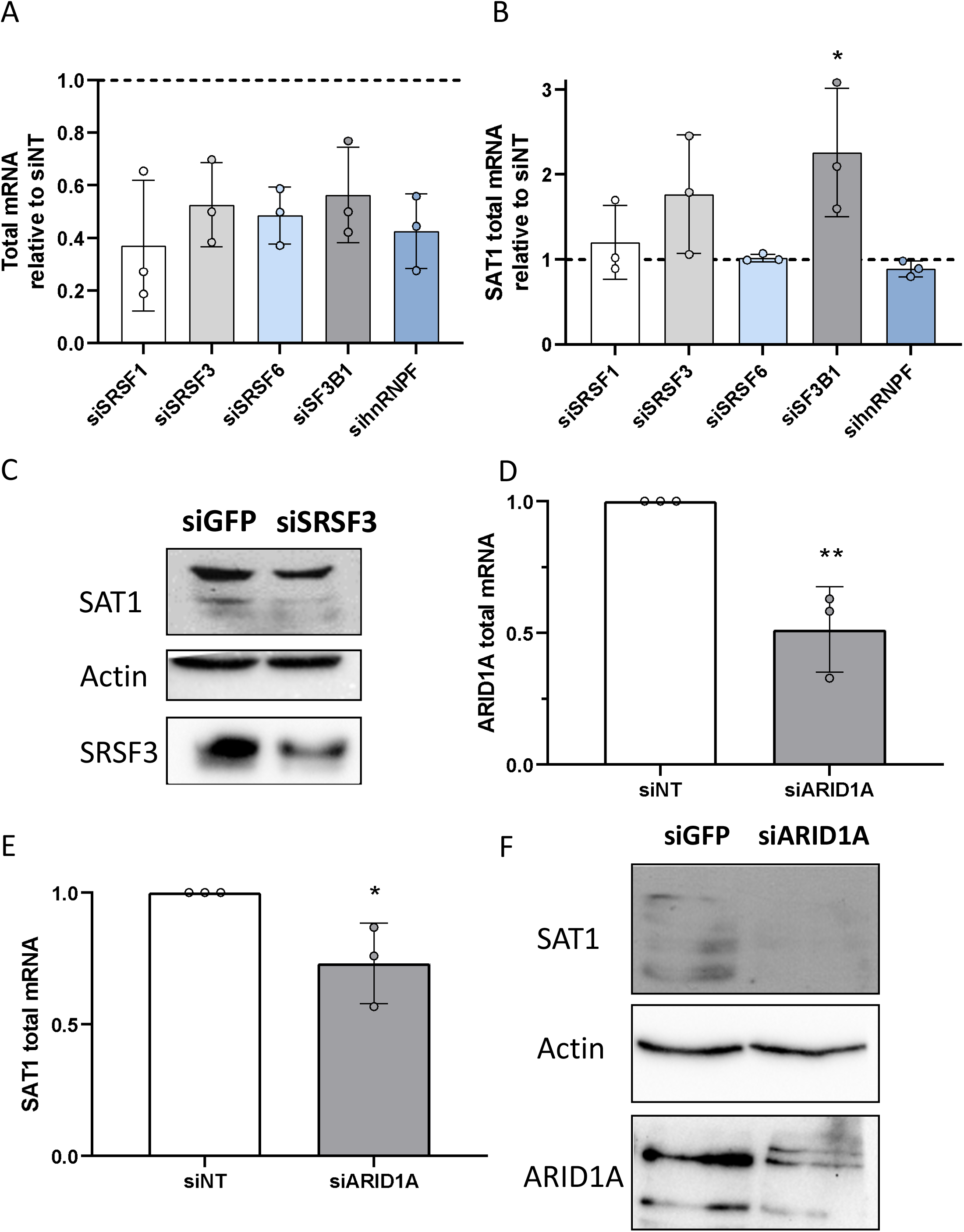
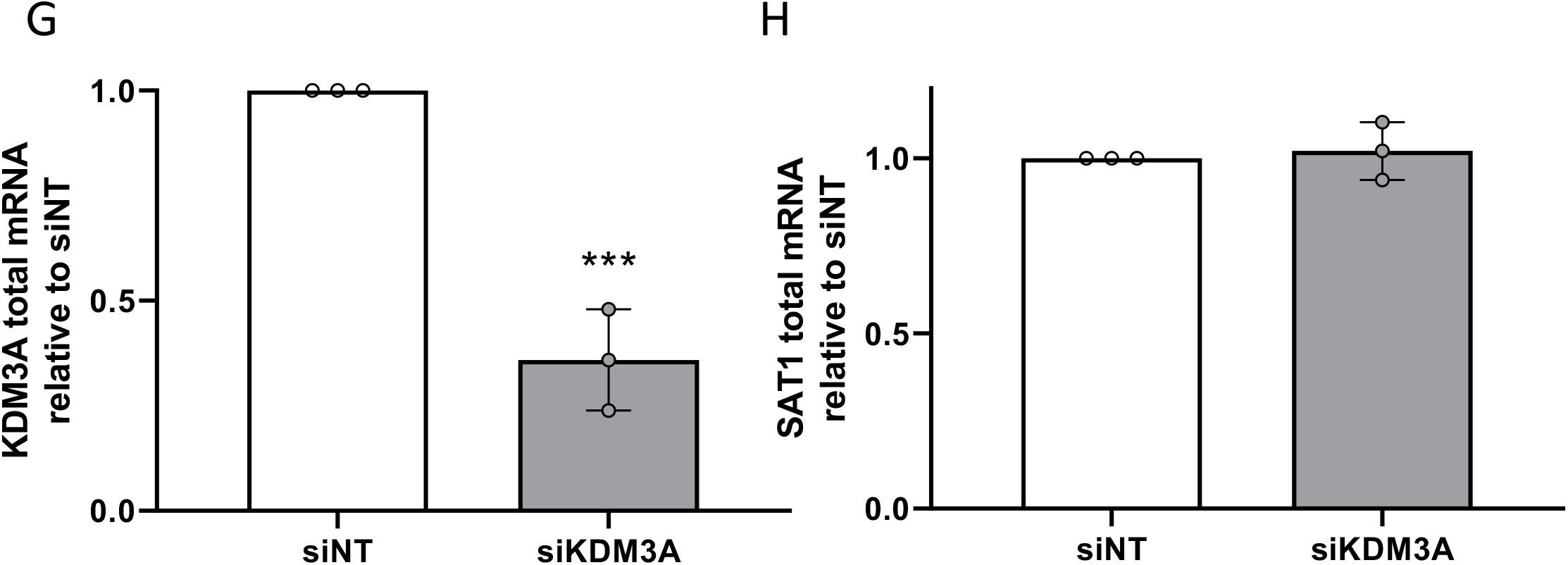
**A-C**. MCF7 cells were transfected with non-targeting siRNA (siNT) or siSF3B1, siSRSF1/3/6 and sihnRNPF for 72 h. Total RNA was extracted and analyzed by real-time PCR for each gene relative to *CycloA* reference gene (**A**) and SAT1 relative to *CycloA* reference gene (**B**). Values in **A** & **B** represent averages of three independent experiments ±SD (* p<0.05; Student’s t test comparing to siNT). Immunoblotting was conducting using the indicated antibodies (**C**). **D-F**. MCF7 cells were transfected with non-targeting siRNA (siNT) or siARID1A for 72 h. Total RNA was extracted and analyzed by real-time PCR for ARID1A (**D**) and for SAT1 (**E**) relative to *CycloA* reference gene. Values in **D** & **E** represent averages of three independent experiments ±SD (* p<0.05, ** p<0.01; Student’s t test comparing to siNT). Immunoblotting was conducting using the indicated antibodies (**F**). **G&H**. MCF7 cells were transfected with non-targeting siRNA (siNT) or siKDM3A for 72 h. Total RNA was extracted and analyzed by real-time PCR for KDM3A (**G**) and for SAT1 (**H**) relative to *CycloA* reference gene. Values in **G** & **H** represent averages of three independent experiments ±SD (*** p<0.001; Student’s t test comparing to siNT).

**Supplementary table S1.**
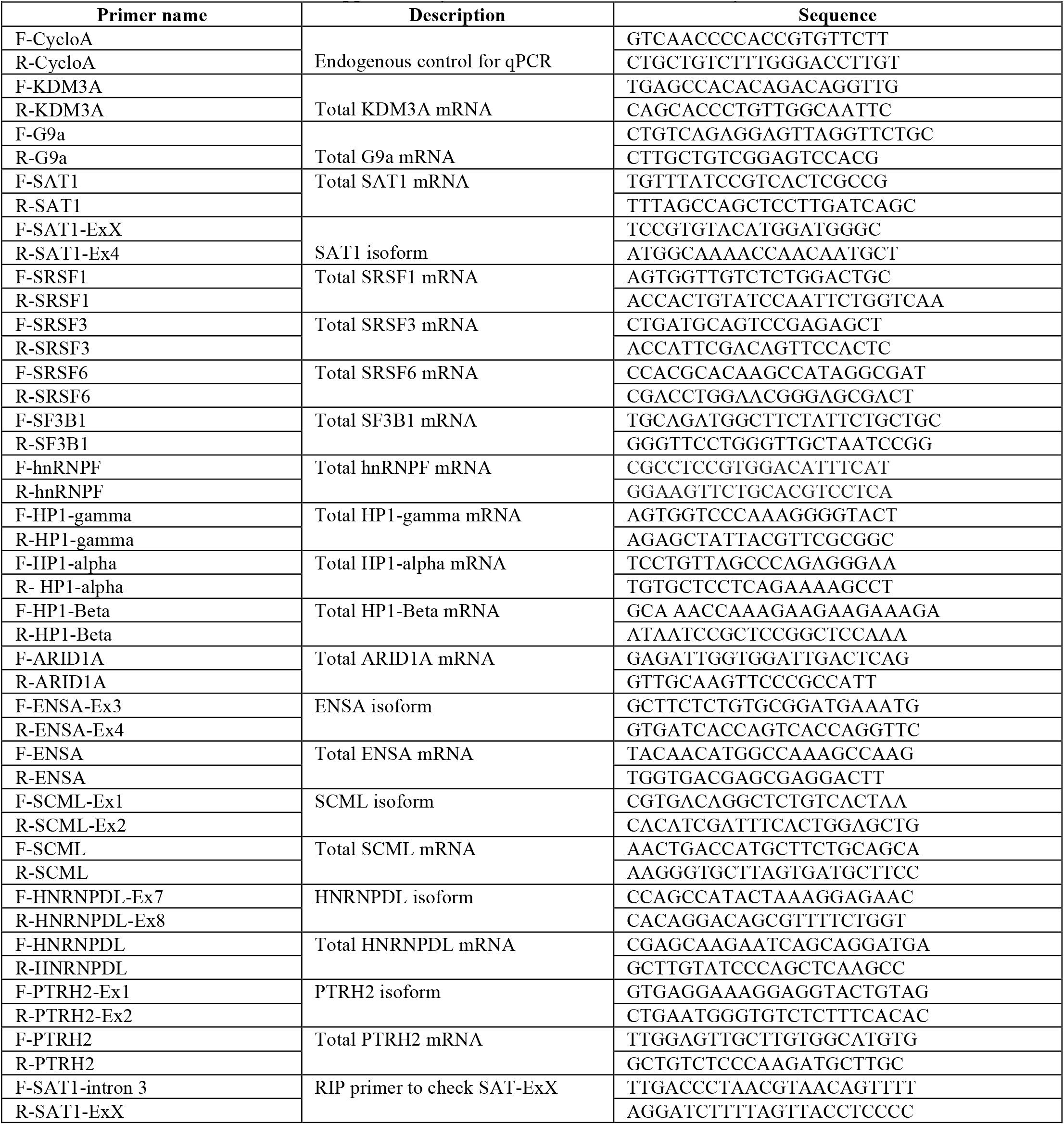
Primer used in this study.

